# Inferring protein ensembles directly from NOESY spectra

**DOI:** 10.64898/2026.08.20.745893

**Authors:** Murray Coles

## Abstract

Solution NMR spectroscopy provides atomistic measurements of proteins in a native-like biophysical state. Because these measurements are ensemble averages, it also has the potential to report on conformational diversity. However, conventional NMR structure determination typically converts experimental observables into restraints for molecular dynamics, which encode information on the mean structure but do not retain information on the underlying conformational distribution. Ensemble selection has long been proposed as an alternative, whereby experimental observables are compared directly with candidate conformers generated independently of the measurements. This allows population distributions to be inferred from the data. However, few such methods have incorporated NOESY - the richest source of structural information in protein NMR - data, due to challenges in the quantitative comparison of experimental and back-calculated spectra. To address this challenge, we previously introduced the CoMAND method, demonstrating that quantitative agreement is practical for NOESY spectra with bespoke heteronuclear editing schemes. Here we extend this approach into a framework for direct inference of protein ensembles within a flexible ensemble-selection architecture incorporating multiple classes of NMR observables. We introduce a quantitative scoring framework for comparing experimental and back-calculated observables and combine it with regularized ensemble selection and Monte Carlo simulated annealing. Integration with the OpenMM molecular dynamics engine allows conformational pools to be generated using established molecular simulation methods. Applied to human ubiquitin, the resulting ensemble provides simultaneous agreement with NOESY, residual dipolar coupling and scalar coupling data while retaining conformational diversity supported by experiment.

## INTRODUCTION

Conformational dynamics are increasingly recognized as contributing to a wide range of protein functions, including catalysis, signaling, transport, and molecular recognition. While protein structure prediction has advanced substantially in recent years, the prediction of such dynamics and their functional roles remains a key frontier ^1–3^. NMR spectroscopy is one of the few techniques capable of providing experimental data in this context, owing to its ability to probe molecules in near-native biophysical conditions - that is, in solution at ambient temperatures and pressures. NMR relaxation measurements can localize motions at an atomistic level and determine their timescales over several orders of magnitude, spanning picoseconds to seconds ^4–6^. Moreover, NMR structural data contain information on the microstates contributing to conformational ensembles and can be used to estimate their populations ^7,8^. This follows from the fact that NMR observables are inherently ensemble averages over molecular conformations; consequently, structural models can only accurately reproduce experimental data if they capture the underlying conformational diversity. However, conventional NMR structures rarely provide this level of detail. This limitation largely arises from the common practice of converting NMR observables into restraints for restrained molecular dynamics (MD). Although restraints derived from ensemble-averaged observables may faithfully represent the experimental data, they do not uniquely specify the underlying conformational distribution. The resulting structures are therefore best described as *uncertainty ensembles* ^8^, where the observed variability reflects uncertainty in interpretation of the restraints, rather than a description of the conformational ensemble giving rise to the data.

Since its inception, biomolecular NMR structures have almost exclusively been determined using restrained molecular dynamics, owing to its ability to search conformational space in the face of intrinsically ambiguous NMR data. The richest form of NMR structural data are NOESY spectra, which report inter-proton contacts at distances of up to ∼6 Å. However, these contacts are identified via cross-peak positions in highly crowded ¹H dimensions. Editing through the attached heavy atom (¹³C or ¹⁵N) to create additional dimensions greatly alleviates this problem, but considerable ambiguity remains. The conventional solution has been an iterative bootstrapping strategy, in which a small set of restraints derived from unambiguous peaks is used to generate initial models, which then serve as filters for identifying additional restraints. The routines for NMR structure determination that have dominated the field for the past 25 years have formalized this approach: restrained molecular simulations tailored for efficient conformational exploration, coupled with algorithms for assigning ambiguous cross-peaks based on intermediate structural models ^9,10^. This can be viewed as a mapping from experimental observation space into a derived restraint space, where the downstream optimization problem is far more tractable. Although these methods have been highly successful, the assignment step inevitably introduces a layer of interpretation, such that this restraint space represents a reduced description of the original spectral information. With the recent breakthroughs in predictive modeling, however, highly accurate protein structures are now almost instantly available ^11,12^, greatly reducing the need to search fold space - and with it reliance on bootstrapping. This opens the way to structure determination protocols that operate directly in observation space, exploiting the full information content of the spectra.

For many years, alternative approaches to constructing conformational ensembles have been proposed, centered on methods that couple experimental data with molecular simulation ^7,8,13–16^. For many folded proteins, where reliable structural models are now available, the problem shifts from determining global folds toward assessing the extent to which experimental data justify deviations from the model used as a structural prior. In this regime, reweighting-based approaches, including ensemble selection (sample-and-select), are particularly well suited ^8,13,15,17^. These methods begin from a pre-generated structural pool and determine populations that reproduce the experimental observables (Fig. 1). This separates the problem of conformational sampling from that of population estimation. Importantly, when formulated within an appropriate uncertainty model, reweighting introduces additional conformational states only when required by the experimental data.

**Figure 1.**
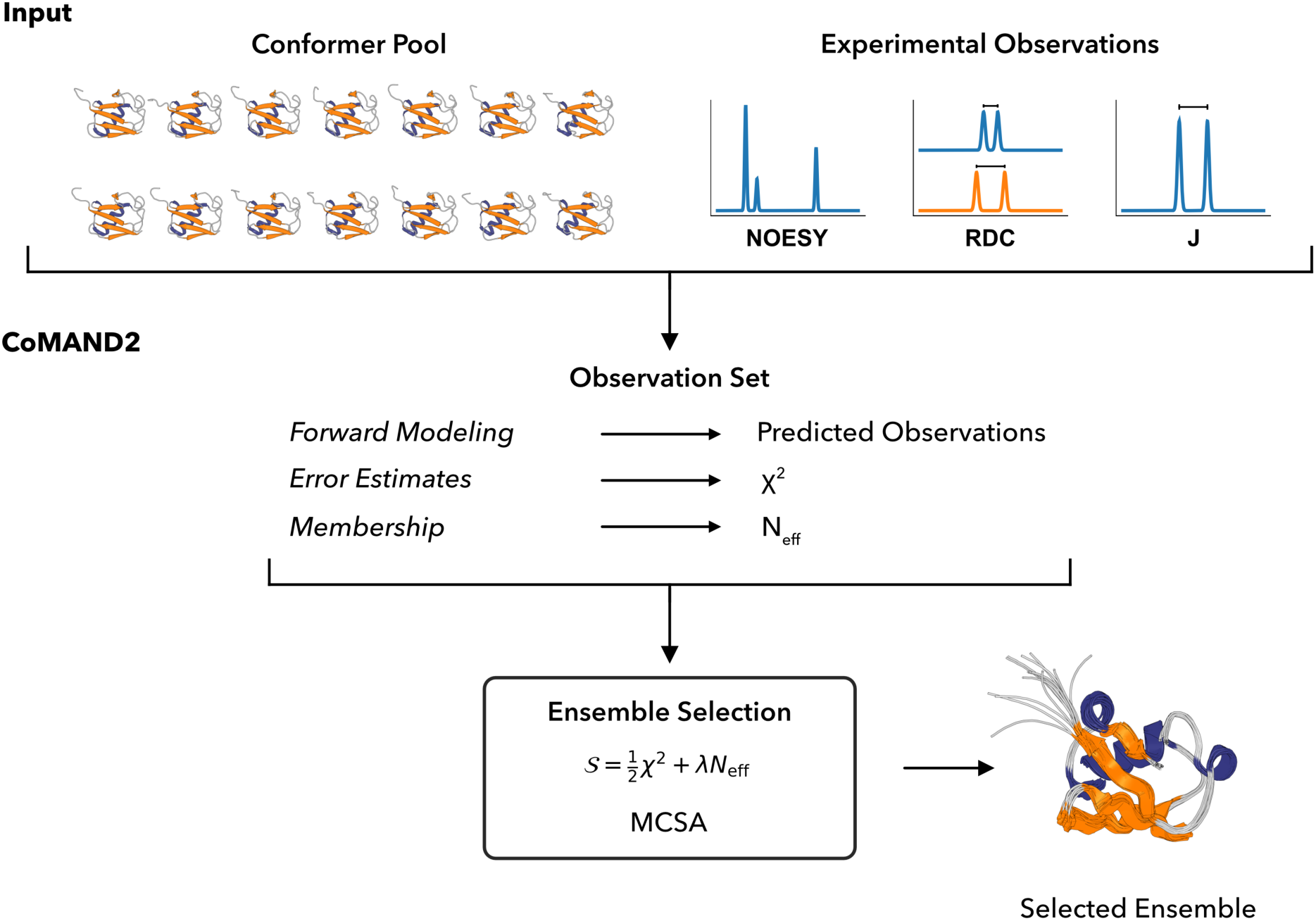
Structure determination using CoMAND2. The workflow is centered around the observation set, which integrates forward models for predicting observables from the conformer pool with error estimates for comparison to experimental data. The modular design allows observation sets containing different experimental data types to be flexibly combined. The scoring function combines weighted contributions from each observation set (*χ*^2^) with a regularization term that reflects ensemble diversity (N_eff_). The final ensemble is selected from the conformer pool by minimizing this score using Monte Carlo simulated annealing (MCSA).

Reweighting approaches rely on methods that allow experimental observables to be back-calculated from conformers in the structural pool. Such methods - referred to as forward models - predict observables from molecular conformations and provide the essential link between simulation and experiment. However, reweighting approaches have rarely incorporated NOESY data as quantitative observables. Although forward models for NOESY spectra have long been available ^18,19^, their integration into ensemble inference has been limited by the absence of robust statistical frameworks for quantitatively comparing experimental and back-calculated spectra. We have shown that such a measure can be obtained using a specific NOESY implementation: the 3D CNH-NOESY experiment ^20^. In this experiment, NOESY correlations are resolved in a ¹³C dimension, exploiting improved chemical shift dispersion and more uniform line shapes, while avoiding the complications associated with indirect ¹H dimensions ^21^. These features enable direct, quantitative comparison between experimental and back-calculated spectra, providing a prerequisite for their incorporation into ensemble selection schemes.

Here, we describe CoMAND2, a platform for protein structure determination via ensemble selection, enabled by quantitative comparison of CNH-NOESY spectra. We introduce a scoring scheme that combines an uncertainty-weighted χ² measure of fit quality with a regularization term penalizing ensemble complexity, enabling explicit control over the trade-off between the two. While the method is centered on NOESY data, it is readily extensible to any experimental observables for which forward models exist. The current implementation also incorporates residual dipolar couplings (RDCs) and scalar couplings. RDCs report on the orientations of inter-nuclear vectors relative to an alignment frame and provide a complementary, global constraint alongside the short-range distance information encoded in NOESY spectra ^22,23^. The resulting ensembles recapitulate all input data and provide a detailed, quantitatively grounded view of the conformers contributing to protein dynamics.

## RESULTS

### The CoMAND2 method

We originally named our method CoMAND - Conformational Mapping by Analytical NOESY Decomposition - reflecting its ability to estimate the populations of individual conformers across defined conformational spaces ^21^ For example, we applied CoMAND to calculate conformational energy maps (cmaps) for use in molecular dynamics protocols. Here, we present a new Python API with the broader objective of providing a platform for NMR structure determination by ensemble selection. The emphasis is shifted to accommodating different classes of NMR observables in a statistically sound manner. The framework integrates three essential components: (i) a molecular representation, (ii) forward models for back-calculating NMR observables, and (iii) quantitative methods for evaluating agreement with experiment.

To represent the molecular topology, we use a conventional chain–residue–atom hierarchy. Atoms retain bonding information and carry NMR-specific attributes required for back-calculation of observables. This system allows specialized external programs to handle reading and manipulation of coordinates, plus deal with exceptions such as post-translational modifications, non-canonical amino-acids, ligands, metals and prosthetic groups. The topology is compatible with any CHARMM-based system, but has a dedicated interface for the openMM molecular dynamics system ^24^.

NMR is unique in providing different types of structural data, each reporting on distinct aspects of biomolecular structure and requiring its own back-calculation method. In our original implementation, we developed the program SHINE, which allows back-calculation of NOESY spectra with any combination of ^1^H, ^13^C and ^15^N dimensions. SHINE applies a standard algorithm based on exponentiation of a relaxation matrix calculated from inter-proton distances (Materials and Methods). The most computationally expensive step is this process is eigen-decomposition of the relaxation matrix, which scales cubically with matrix size. To reduce this cost, we adopt a fragmentation approach, calculating matrices for subsets of the structure surrounding each atom of interest. This provides a very close approximation of the full relaxation matrix at a fraction of the computational cost. For 3D CNH-NOESY spectra, fragments are centered on a single ^15^N-bound proton. The back-calculated spectrum for each fragment is thus a 1D vector (“strip”) along the ^13^C dimension (Fig. 2). These strips are then compared to the corresponding vectors extracted from the experimental spectrum.

**Figure 2.**
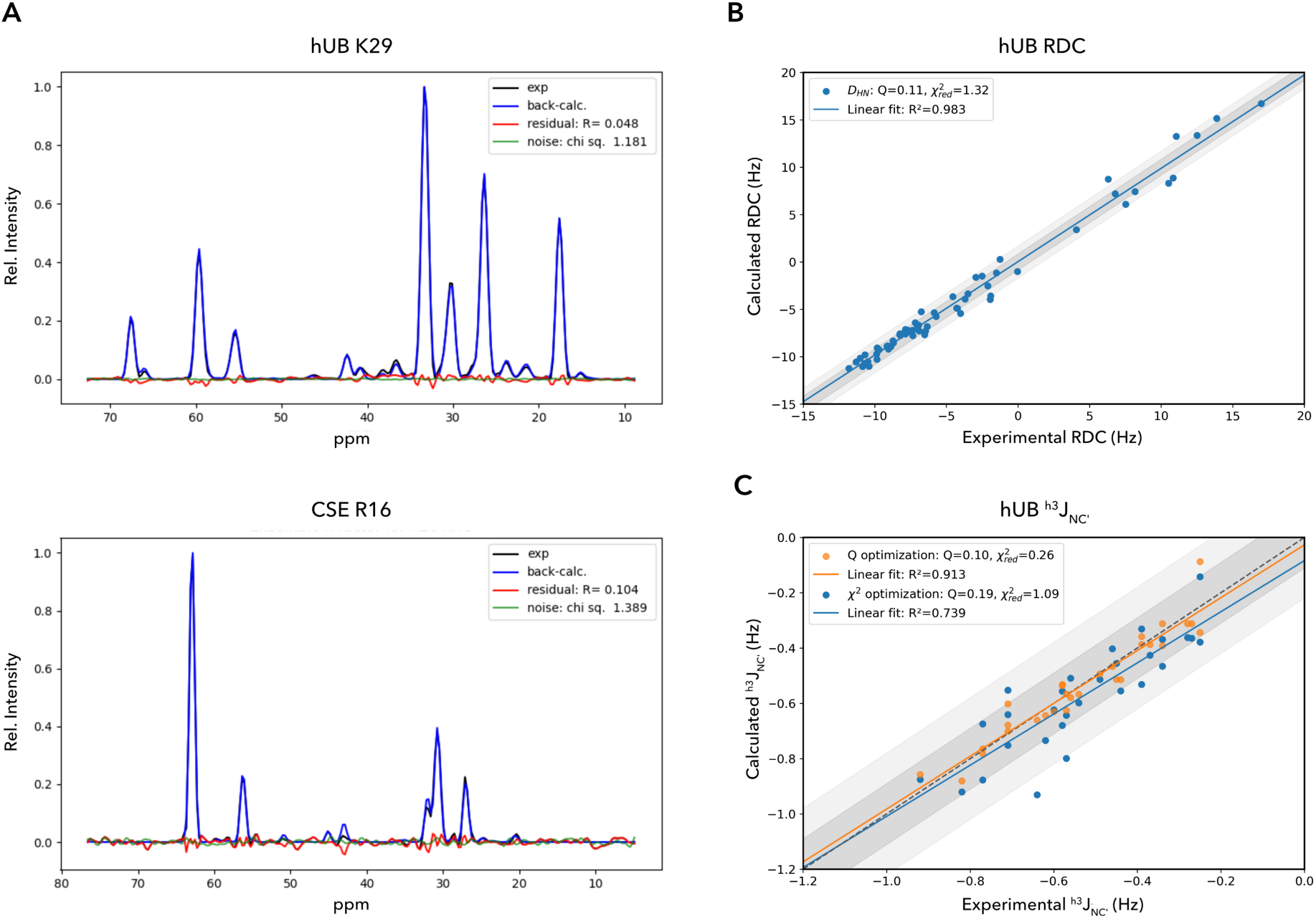
Examples of the scoring function applied to three classes of NMR data. (A) Ensemble selection fitting of individual strips extracted from 3D CNH-NOESY spectra of hUb (top) and the small calcium-sensing protein CSE ^40^ (bottom). Experimental data are shown in black, back-calculated strips in blue, residuals in red, and simulated noise traces generated from the corresponding noise models in green. The noise models were calibrated by sampling empty spectral regions and define the baseline component of the heteroscedastic noise model used for 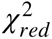 calculation (Materials and Methods). (B) Ensemble selection fitting of RDC data for hUb. Ensemble selection was performed using a dataset containing multiple coupling types measured in a single alignment medium (5% 3:1 DMPC:DHPC neutral bicelles). The panel shows the correlation between experimental and back-calculated D_HN_ couplings. A linear regression is shown in blue, with the identity line (y=x) shown as a grey dashed line. Shaded regions indicate deviations of 1σand 2σ, where σ(0.85 Hz) was estimated from the residual distribution and used for calculation of the 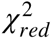 value (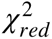= 1.32). (C) Ensemble selection of ^h3^J_NC’_ couplings across hydrogen bonds for hUb ^33^ using two optimization objective functions. Q-factor optimization (orange) minimizes the normalized residual without reference to experimental uncertainty, whereas the χ² objective (blue) includes a minimum contribution, 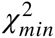= 1, to prevent fitting below the expected noise level. Regression lines and shading are as for panel B, with σ= 0.12 Hz. Evaluation of the ensemble optimized by Q-factor gives 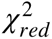 = 0.32, indicating fitting beyond the level justified by the experimental uncertainty.

For RDC and scalar coupling data, we have adopted established methods for back-calculation from coordinates (Materials and Methods). To integrate these data with the NOESY framework, we have implemented an extensible system based on the concept of *Observations*: 1D vectors of experimentally related values. For example, an observation could be a 1D strip extracted from a NOESY spectrum or a list of RDCs measured in the same alignment medium. Observations are initialized with the experimental data and a method for evaluating the agreement with back-calculated data. These are collected into *ObservationSets* that are initialized with the back-calculation method and parameters. This system allows NMR data to be flexibly grouped and weighted in downstream ensemble selection procedures.

### Scoring and Regularization

Any refinement procedure requires a quantitative measure of agreement between experiment and prediction. More fundamentally, it requires a definition of the level of disagreement that can reasonably be attributed to experimental and forward-model uncertainty, since only residual discrepancies beyond this level provide evidence for deficiencies in the structural model. In the original implementation of CoMAND, this agreement was quantified using an R-factor based on the relative root-mean-square deviation (RMSD) between experimental and back-calculated NOESY spectra. The normalization by the total signal magnitude makes the metric closely analogous to the Q-factor used for RDCs ^25^ and the R-factor employed in X-ray crystallography. However, when combining heterogeneous experimental observables, such relative measures of fit quality are insufficient, as they provides no objective criterion for distinguishing expected discrepancies from those that indicate deficiencies in the structural model (Fig. 2C). Instead, the scoring function must account for the effective uncertainty associated with each observable, encompassing both experimental error and limitations of the forward model. Such uncertainty-weighted residuals arise naturally from the likelihood function in Bayesian inference and maximum-entropy ensemble refinement, where they define the boundary between agreement attributable to noise and residual disagreement that reflects missing structural detail. We have therefore adopted the reduced χ² statistic for optimization. For a single observation vector, *j*, it evaluates the deviation of each data point, *i*, relative to an estimate of the variance:

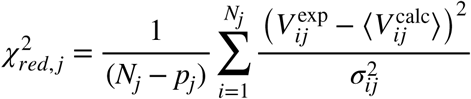

where 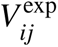 is the experimental value, 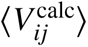 is the ensemble average of the back-calculated values and 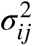 is the variance, estimated from the measurement error. The reduction factor, 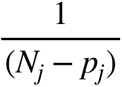, reflects the degrees of freedom, where *N_j_* is the number of observations and *p_j_* the number of fitted parameters, which depends on the data type (Materials and Methods).

The χ² scoring function requires an estimate of the variance associated with each data point along the observation vector. These estimates are a critical component in integrating data from multiple sources, as they determine the relative statistical weight assigned to each observable. Underestimation over-emphasizes the corresponding data type and promotes absorption of forward-model error into the structural model, whereas overestimation diminishes its contribution and reduces sensitivity to genuine structural information. For scalar couplings, experimental uncertainties generally dominate the overall forward-model error, and point-wise variance estimates derived from experimental measurements are adequate. For RDCs, however, uncertainties in the forward modeling cannot be neglected, arising for example from the alignment description and physical assumptions, such as fixed bond lengths. Accordingly, experimental errors for RDCs should be regarded as lower bounds on the effective variance. Several approaches to obtaining more realistic estimates have been explored in the literature. In the present work, we adopt an iterative estimator based on residual deviations that captures an effective contribution from forward-model uncertainty while remaining deliberately conservative to avoid over-interpreting residual noise as conformational heterogeneity (Fig. 2B).

To accommodate NOESY data within the χ² scoring framework, the effective variance, 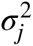 is modeled as the sum of a baseline contribution and an intensity-dependent term:

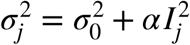

where 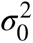 is the baseline variance and *I_j_* denotes the total absolute signal intensity for strip *j*. This model accounts for both the heteroscedastic experimental noise, including t₁ noise, and the dominant intensity-dependent contributions to the forward-model error, such as mismatch between experimental and simulated line shapes. To estimate 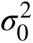, we construct a baseline noise model by sampling empty regions of the experimental spectrum. The parameter α is then determined from the dependence of the overall variance on intensity for a selection of well-described strips (Fig. S1). In conventional NOESY spectra, the total intensity is typically dominated by the diagonal signal, whose intensity often exceeds the combined intensity of all other peaks. As a result, estimates of the intensity-dependent term are biased by contributions from structurally uninformative signals. The 3D-CNH-NOESY experiment has a significant advantage in that it intrinsically lacks diagonal signals, allowing the error estimate to more directly reflect the uncertainty associated with cross-peaks. Consequently, the fitted parameter αis small relative to the baseline contribution, providing a robust variance estimate for χ² calculations. For most conventional NOESY spectra, the intensity-dependent term would amount to a major contribution to total noise, as intensity is dominated by the diagonal signal, which will often be far larger than the sum of all other peaks. However, the 3D-CNH-NOESY has a significant advantage in that it intrinsically lacks diagonal signals. This allows αto be small, yielding a robust estimate of the variance for χ^2^ calculations.

Although χ^2^ provides the natural objective function for ensemble selection by expressing deviations relative to the experimental uncertainty, it does not characterize the scale of the experimental signal. In this respect, the R-factor - and the analogous Q-factor for RDC data - remains informative as a companion metric. Whereas χ^2^ quantifies agreement relative to the noise level, the R-factor expresses the same residuals relative to the magnitude of the experimental signal. Together, these distinguish the quality of fit from the quality of the underlying dataset.

Another important aspect of scoring in ensemble selection is model complexity. For an ensemble representation, there is potentially one additional fitting parameter per distinct conformer in the ensemble, corresponding to its statistical weight, subject to a normalization constraint. Each additional weight increases the dimensionality of the model and reduces the effective degrees of freedom in the χ² statistic. If this is not properly controlled, larger and more heterogeneous ensembles will systematically achieve lower χ² values simply by absorbing experimental noise; i.e. overfitting. It is therefore essential to quantify model complexity explicitly. To this end, we define an effective ensemble size, N_eff_, which reflects the number of conformers contributing to the ensemble. The conformational space is first subdivided into states. These can be discrete - e.g., side chain rotamer bins - or based on continuous probabilistic distributions. A membership matrix can then be defined with elements giving the probability that conformer *j* belongs to state α. The N_eff_ score is defined as the Hill number of order 2 (inverse Simpson index) computed over state occupancies ^26^:

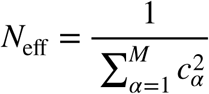

where *M* is the number of states and *c_α_* denotes the normalized occupancy of state *α*obtained by summing membership weights over all structures in the ensemble (Materials and Methods). Because N_eff_ depends quadratically on the populations, it favors compact population distributions over solutions in which population is dispersed across many weakly populated conformers ^27^. The minimum value of N_eff_ is 1, corresponding to the case in which all ensemble weight is concentrated in a single state, whereas larger values arise as membership probabilities become more evenly distributed. N_eff_ therefore acts as an entropy-based regularization term, reflecting model dimensionality. A scoring term combining χ² with N_eff_ thus provides an explicit trade-off between residual error and model complexity:

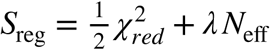

where λis a regularization factor that controls the balance between the two and is estimated by analyzing the response of χ² and N_eff_ to small perturbations around an initial solution (Materials and Methods). In this form, the scoring function is analogous to the Akaike Information Criterion (AIC), with N_eff_ acting as a continuous proxy for the number of model parameters.

In ensemble selection approaches, the candidate pool defines the hypothesis space over which population inference is performed. Implicit in its construction is a partitioning of conformational space into states that can be distinguished by the experimental data. Likewise, the state space used to define N_eff_ should reflect aspects of conformational complexity that the experimental data are capable of resolving. We have previously shown that CNH-NOESY observables are highly sensitive to local dihedral geometry ^21^. Accordingly, an effective regularization space for residue *r* comprises the three backbone dihedrals (ψ_r-1_, ω_r-1_, and φ_r_) and up to two side-chain dihedrals (χ_1_, χ_2_). In the present implementation, the partitioning of this space is defined by Gaussian mixture models fitted to empirical dihedral angle distributions, ensuring that all states represent physically realistic conformers (Fig. 3). This residue-level formulation naturally decomposes the regularized objective function into independent, residue-level contributions. Other regularization spaces can be defined for specific applications, provided they reflect conformational distinctions that are meaningful for the inference problem. Figure 5 illustrates this flexibility, using hard-membership to partition side-chain rotamer space.

**Figure 3.**
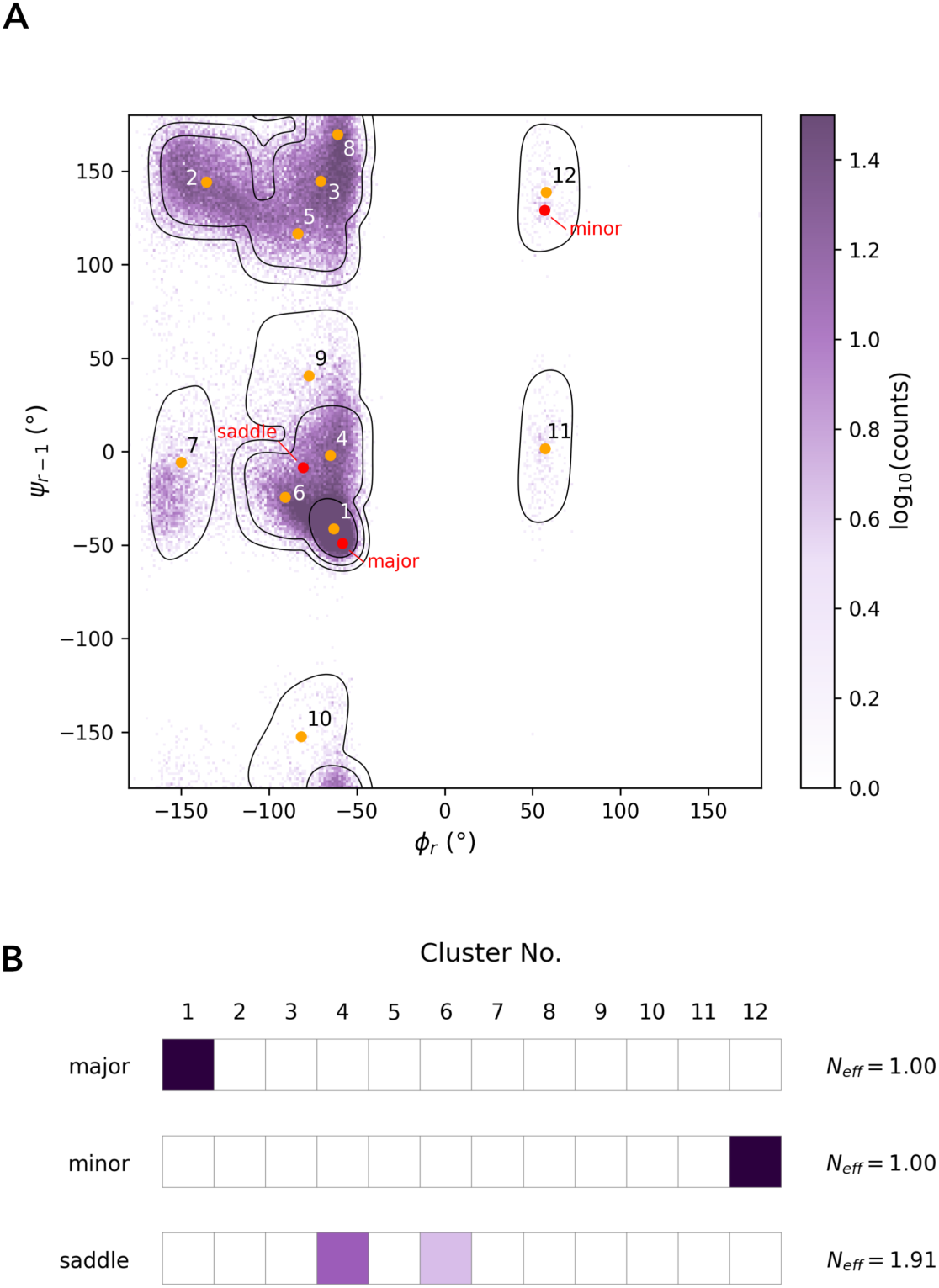
Regularization using soft membership. An example of the soft membership space used for regularization is shown for alanine. (A) The density distribution of ∼150,000 alanine residues from the MolProbity database ^38^ is shown in the (ψ_r-1_, φ_r_) sub-space. Contours represent the probability density of a Gaussian mixture model fitted to the data, drawn at equally spaced intervals of log_10_(p), with the lowest contour at log_10_(p) = -1. Cluster centers are shown as orange circles and numbered according to their mixture component weight. The membership vector for any conformer is calculated from the probability of assigning that conformer to each cluster. Membership vectors for the three points highlighted in red are shown in panel B. Note that that the space presented is a two-dimensional projection of the full (ψ_r-1_, ω_r-1_, φ_r_) space used for alanine and that fitting is performed on a unit vector representation of each dihedral. (B) Membership vectors for the three conformers highlighted in panel A are shown as color-coded probability bars, with values from 0 to 1 represented from white to dark purple. N_eff_ values are shown for each example. N_eff_ reflects ambiguity in state assignment rather than state probability, consequently conformers from basins with both high and low populations can have values approaching 1. In contrast, conformers located between highly populated basins exhibit distributed membership and therefore higher N_eff_.

### Ensemble Selection and Optimization

The scoring system described above provides the objective function for ensemble selection. For each observation, the ensemble average is assumed to be a linear combination of the observations for its individual members, an approximation justified when the lifetimes of conformational states are long relative to their interconversion times. Since each back-calculated observation is a vector, all observations for the pool can be compiled into a matrix, and the ensemble average is obtained by multiplying this matrix by a vector of weighting factors. The general scheme of reweighing methods involves evolving these weighting factors to minimize the scoring function. In the ensemble selection method applied here, the weights are integers indicating how often a member of the pool is included in the ensemble; capping the weights at one enforces a unique ensemble in which each pool structure appears at most once.

In the original CoMAND protocol, we implemented a greedy (steepest descent) algorithm for ensemble selection. At each iteration, every conformer in the pool is evaluated, and the one that most reduces the overall score is added to the ensemble. While deterministic, this algorithm does not produce ensembles of a fixed size, proceeding until no further score reduction is possible. Moreover, the scoring functions defined here - including data from several sources and including a N_eff_ penalty for model complexity - can represent a more rugged scoring landscape where the greedy routine can become trapped in local minima. To address this limitation, we extend the framework with a Monte-Carlo Simulated Annealing (MCSA) algorithm, following the strategy introduced by Chen *et al.*^17^ and subsequently applied in the Al Hashimi group ^15,28^. The MCSA routine begins with an arbitrary ensemble and randomly swaps one member for a pool member at each iteration. Changes that improve the score are accepted under a standard Metropolis simulated annealing criterion (Materials and Methods). This procedure is stochastic, allowing repeated runs to assess the consistency of results. It also naturally accommodates ensembles of a fixed size, a requirement for downstream optimization protocols.

A potential limitation of ensemble selection approaches is the conformational diversity of the structural pool: conformations can only contribute to the final ensemble if they are represented in the pool. One way to address this is through targeted refinement, for example by using the residue-based scoring scheme to carry out optimization at the level of individual residues or short peptides. In such optimizations, the minimum achievable score is often limited by the quality of the conformer pool. In most cases, poorly explained residues arise from inaccuracies in the starting models and can be remedied by local rebuilding. It is also possible that the pool does not adequately sample conformers needed to describe the data. This is particularly true for backbone polymorphisms, where a short peptide segment adopts multiple distinct conformations (Fig. 4). In these situations, it is important to ensure that the structural pool maintains sufficient diversity around each conformer to allow the data to be accurately reproduced.

**Figure 4.**
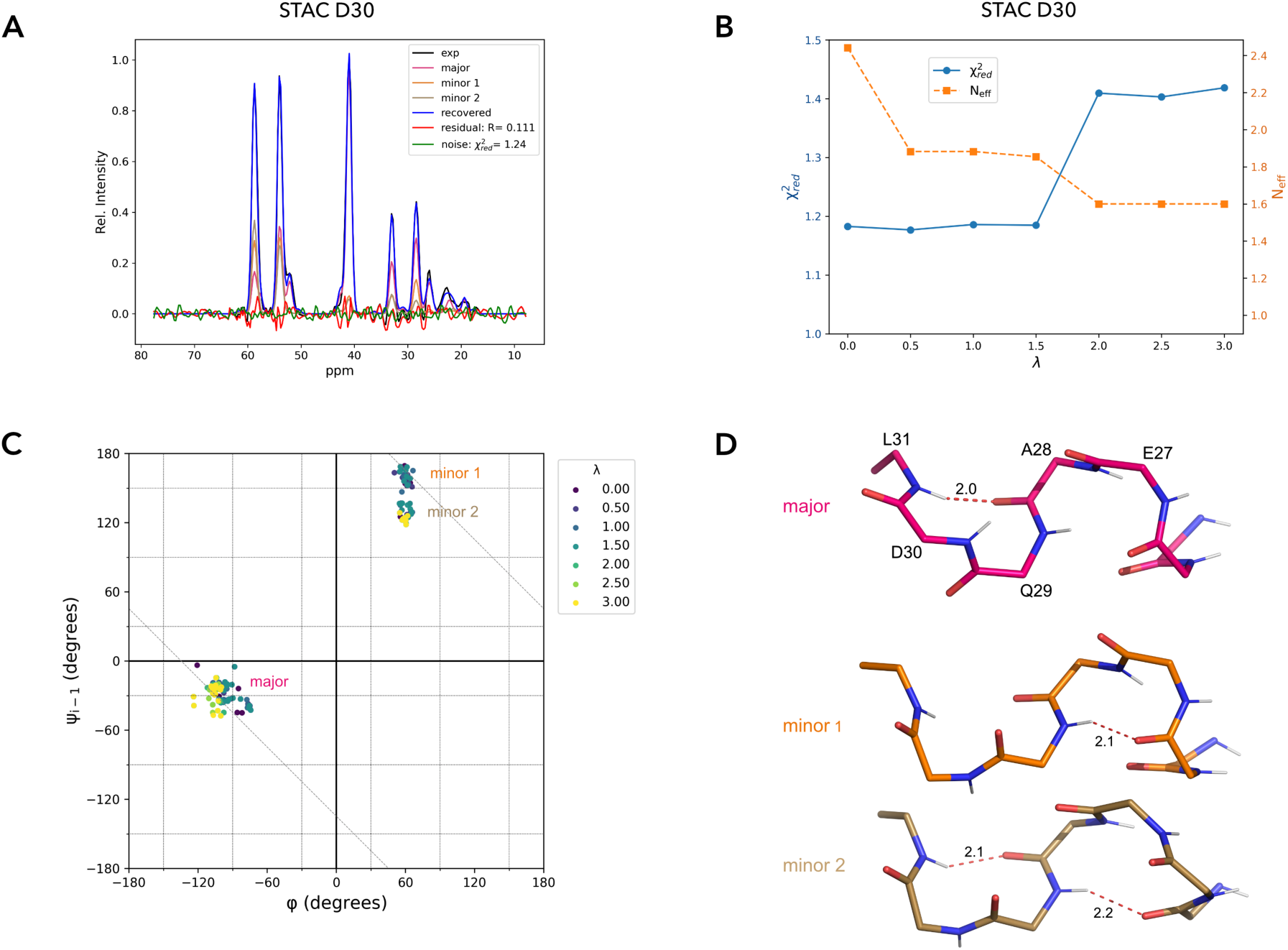
Regularization preserves an experimentally supported backbone polymorphism. The effect of regularization on ensemble selection is shown for a backbone polymorphism identified in the isolated STAC domain protein Af1502 ^29^. (A) The CNH-NOESY data is not well described by a single backbone conformation. In order to describe the data well, the back-calculated strip recovered by ensemble selection (blue) requires contributions from three distinct conformers (See also Fig. S2). (B) Dependence of the effective ensemble size (N_eff_) and 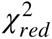 on the regularization parameter λ_Neff_. (C) Corresponding redistribution of the ensemble in backbone conformational space, showing persistence of the major and minor conformational states. The minor state contains an additional substructure that collapses only at higher regularization strength, accompanied by a marked increase in 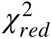. All three conformational states are therefore supported by the data. The dashed diagonal line linking the major and minor conformations indicates that they can be inter-converted by rotation of the plane of the Q29-D30 peptide bond. (D) Representative structures of the backbone conformational states retained after regularization (λ_Neff_ = 0.25). The major conformation (62%) is shown in purple, while the minor conformations (∼19% each) are in dark and light orange. Hydrogen bonds are shown in red and labelled by acceptor-donor distance (Å). Flipping of the Q29–D30 peptide bond alters the hydrogen-bonding pattern: the major and minor 2 conformations consistently form an L31–A28 hydrogen bond, whereas this interaction is absent in minor 1. Both minor conformations favor retention of the canonical hydrogen-bonding pattern of the preceding helix.

The accuracy of ensembles obtained by selection depends critically on the quality of the conformer pool. This suggests that further refinement can be achieved by iteratively evolving its contents. A key advantage of ensemble selection methods is that they are largely agnostic to the source of conformers, such that a wide range of sampling strategies can be employed for this task. Unrestrained molecular dynamics is particularly attractive, as the resulting conformers are thermally consistent and connected by physically plausible interconversion pathways. Here we introduce a replica-exchange MD protocol for refinement, in which multiple replicas undergo short cycles of simulated annealing. Each cycle contributes new conformers to the pool, from which an updated ensemble is selected using the MCSA procedure. If the resulting ensemble improves the overall score, it is used as the starting point for subsequent sampling. Individual conformers are implicitly graded by their frequency of selection, and those that are consistently under-represented are replaced, once a predefined pool size is reached. In this way, the pool becomes progressively enriched in conformers required to explain the data. An additional advantage of this framework is its flexibility: sampling conditions can be adjusted during the refinement process, for example by introducing restraints or biasing potentials at early stages. Figure 6 shows an example of this refinement protocol for human Ubiquitin.

### Examples: regularization in continuous and discrete spaces

The STAC domain is a small four-helix bundle associated with prokaryotic signal transduction, occurring either as an independent protein or as a domain of transmembrane solute-carrier systems ^29^. During structural analysis of Af1502, a single STAC-domain protein from *Archaeoglobus fulgidus*, CoMAND analysis suggested backbone polymorphism in the α2-helix capping motif, as the observed CNH-NOESY data for D30 could not be adequately explained by a single conformer (Fig. 4 and Fig. S2). Under the current implementation, this hypothesis can be examined directly by constructing a regularization curve, i.e. a plot of 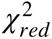 as a function of λ_Neff_, which captures the trade-off between agreement with the experimental data and conformational complexity (Fig. 4B). To construct this curve, we performed ensemble selection using the MCSA protocol, optimizing the regularized scoring function in the default dihedral-based regularization space described above. At each value of λ_Neff_, we selected ensembles of 8 members from a pool of 1260 conformers, pooling the results of 10 independent optimization trials. Across the range of λ_Neff_ values examined, all ten optimization trials converged on the same backbone polymorphism, reproducibly comprising major and minor conformational states related by rotation of the Q29–D30 peptide bond (Fig. 4C). Increasing λ_Neff_ did not eliminate the major conformational states. Instead, collapse of a substate within the minor conformation at λ_Neff_ = 1.5 was accompanied by a substantial increase in 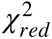 value, indicating that this additional conformational heterogeneity is supported by the experimental data. The three conformational states are distinguished by distinct hydrogen bonding patterns (Fig. 4D).

The system outlined above represents one possible regularization approach; however, bespoke state spaces can also be defined for specific applications. We have previously applied CoMAND analysis to peptides designed as single α-helices, stabilized by glutamine side chain–backbone hydrogen bonds facilitated by an ALQ sequence repeat ^30,31^. Our original analysis of the peptide (P3-7)_3_ suggested an enrichment of glutamine side-chain conformations compatible with hydrogen bond formation, relative to the expected background ^31^. We now test this hypothesis using a discrete rotamer-bin representation of side-chain conformational space (Fig. 5). In such discrete spaces, the N_eff_ score reflects only the number of occupied bins and is therefore insensitive to the distribution within each bin. In contrast to the continuous space described in the previous example, membership probabilities are not weighted by a database prior; all rotamer bins are treated equally. This avoids biasing the inferred rotamer populations toward the background distribution. We constructed a regularization curve for Q14 in the (P3-7)_3_ peptide, selecting from a pool of 26000 frames drawn from MD simulations using two different forcefields ^31^. This shows a rapid collapse toward a single conformer (N_eff_ score ∼1) as the regularization strength increases, at very modest cost in terms of fit quality (Fig. 5A). Thus the data are compatible with a single side chain rotameric state. In line with our hypothesis, regularization also shifts the rotamer population toward states compatible with side chain–backbone hydrogen bonding. These examples demonstrate how the residue-level scoring framework can be flexibly applied to balance experimental fit and conformational complexity across both continuous and discrete state spaces.

**Figure 5.**
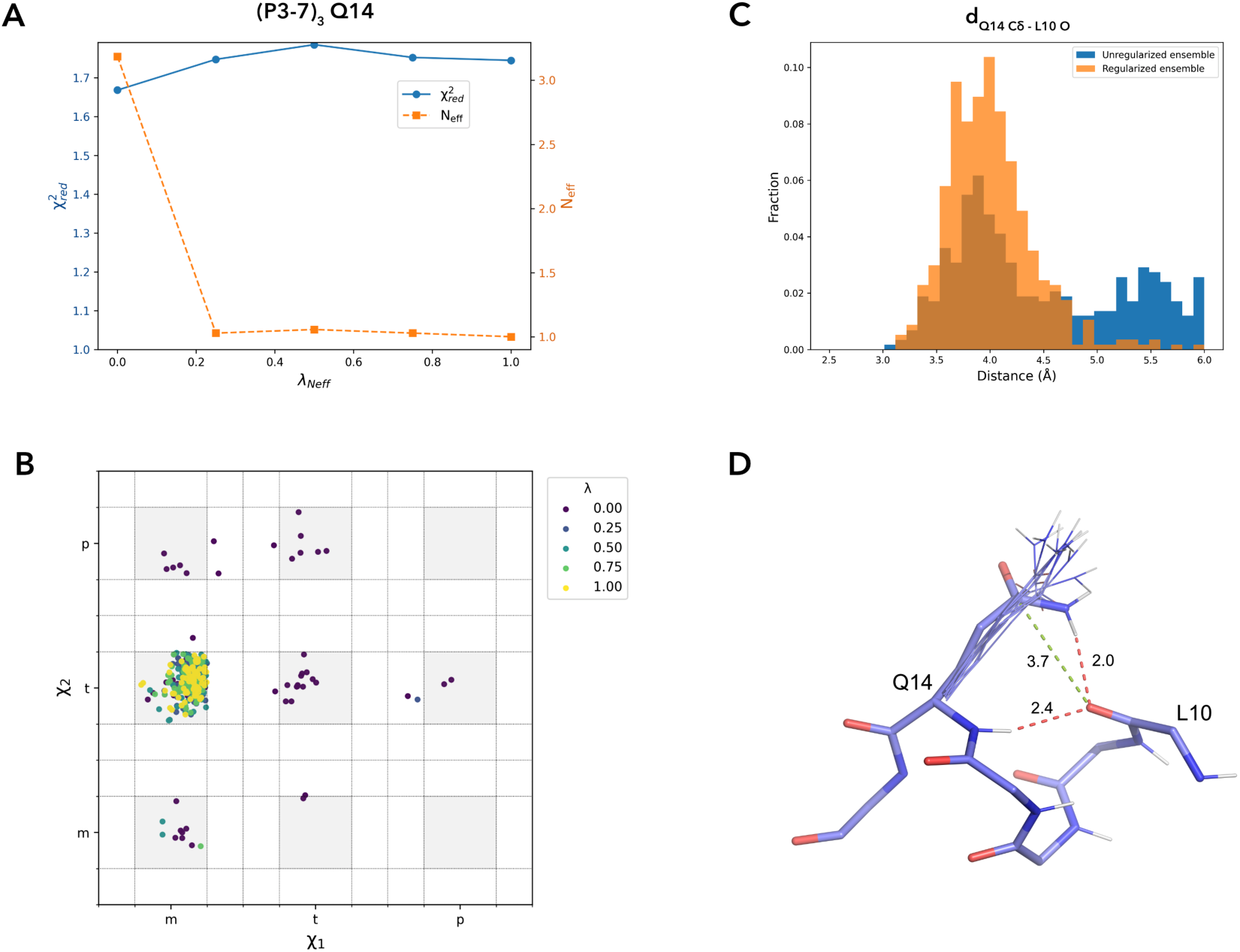
Discrete regularization selects a hydrogen-bond-competent side-chain conformation. The effect of regularization on ensemble selection is shown for Q14 in the single α-helical peptide (P3-7)_3_ ^30,31^. (A) Dependence of the effective ensemble size (N_eff_) and 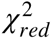 on the regularization parameter λ_Neff_. Increasing λ_Neff_ rapidly selects a single side-chain conformer, with only a modest increase in 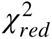 indicating that the selected conformer retains compatibility with the experimental data. (B) Corresponding redistribution of the ensemble in the χ_1_/χ_2_ rotamer space, with population concentrated in the mt rotamer. (C) Distribution of the Q14 C_δ_ –L10 O distance before and after regularization. Regularization enriches conformers with geometries competent to form a side chain–backbone hydrogen bond, increasing the fraction with C_δ_ –O distances below 4.2 Å from 37% to 60%. (D) Representative conformers from the regularized ensemble (λ_Neff_ = 0.25) showing the selected Q14 side-chain orientation. Side chain–backbone hydrogen bonds in these peptides are typically bifurcated with the canonical backbone hydrogen bond. An example of this type of interaction is shown in red, with the C_δ_ –O distance in green.

### Example: ensemble selection for human Ubiquitin

To test the original CoMAND method, we calculated an ensemble for human Ubiquitin (hUb), widely considered the gold standard for NMR structure determination, where a range of reference ensembles are available, compiled under different criteria. Here we update that ensemble using the current scoring and regularization protocol and including literature RDC data for four alignment media ^32^, ^3^J_HNHα_ scalar couplings and ^h3^J_NC’_ couplings across backbone hydrogen bonds ^32,33^. As a conformer pool, we used the same set of 40000 frames drawn from equilibrium MD used previously (Materials and Methods). We first selected an ensemble of 12 structures from this pool using the MCSA method, then applied the replica-exchange MD protocol described above for refinement. The resulting ensemble and breakdown of residue-based scoring is shown in Figure 6. Comparisons to the previous ensemble (6QF8) and three literature ensembles (2MJB, 2NR2 and 2KOX) as shown in the Supplementary Material. The new ensemble for hUb show very good agreement with all classes experimental data. The average 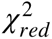 for CNH-NOESY data is 5.7, with a median of 4.2. The average R-factor is 0.14, reflecting the high quality of the data. For RDCs, the average 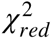 across the four alignment media is 1.40 with a Q-factor of 0.16. For scalar couplings, the equivalent measures are 1.87 and 0.21, respectively. A detailed summary of scores and structure quality measures is provided in Table S1.

**Figure 6.**
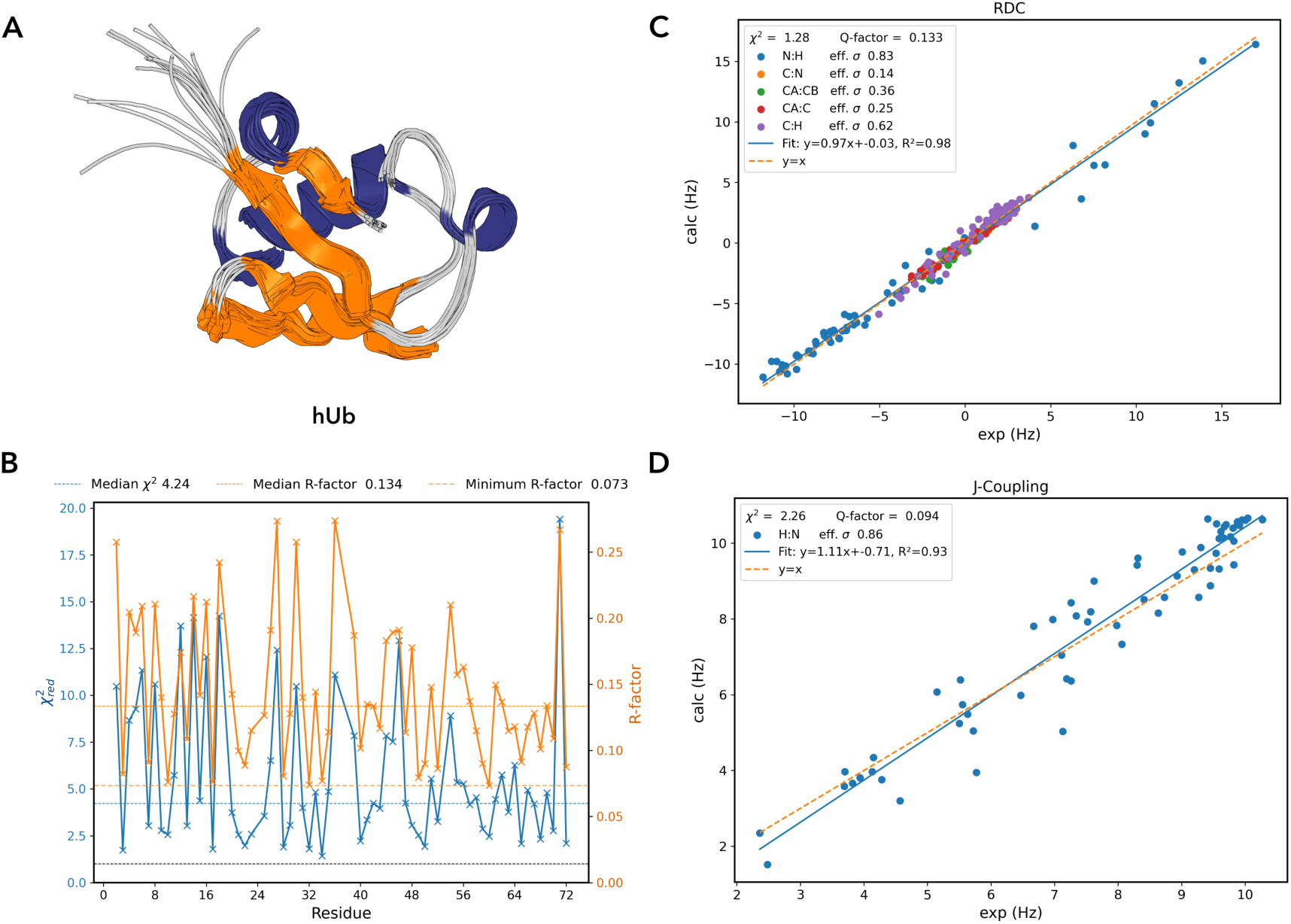
Ensemble selection for human Ubiquitin. (A) The 12-member ensemble selected for hUb using CNH-NOESY, RDC and scalar-coupling data. Structures are superimposed over the backbone of ordered residues (Q2–R72) and shown relative to the average structure (RMSD = 0.56 Å). (B) Residue-level scoring of the CNH-NOESY data for the ordered region of the selected ensemble. Both 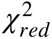 and R-factors are shown for all residues for which data are available. Horizontal lines indicate the median 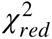 and R-factor values, plus the minimum R-factor. (C) Experimental versus back-calculated RDCs for the Pf1 alignment medium ^32^, with points colored by coupling type. The solid blue line shows a linear regression and the dashed orange line the identity relationship (y=x). Effective standard deviations estimated from the regression are shown for each coupling type and used to calculate 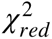 (1.28). (D) Experimental versus back-calculated ^3^J_HNHα_ scalar couplings, shown as in (C).

The value of 5.7 for the CNH-NOESY data may at first appear high in comparison with the other data types, but reflects the very high sensitivity of NOESY to small distance changes. Optimization therefore reaches a plateau over the final few units of S_reg_, with respect to meaningful conformational differences. This is particularly true for the high-quality hUb data, where changes of 1-2 units in 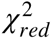 correspond to very small changes in the relative intensities of back-calculated peaks, as reflected in the attendant R-factors. The small difference between the median and minimum R-factor values (0.134 versus 0.073) is therefore informative, indicating that the majority of residues have reached this plateau in the optimization. Nevertheless, the selected ensemble, several residues with higher 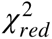 scores remain. For most, acceptable scores can be achieved by local rebuilding, for example by altering side-chain rotamer distributions. For others (e.g. T12, T14, E18 and L71), this is not the case. These residues - prominently featuring β-branched amino acids isoleucine, valine and threonine in either the *i* or *i-1* positions - appear to require conformers that are rare in the structural pool. For example, the 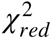 score for L71 is highly sensitive to the side chain conformation of V70, but the required side-chain rotamers are strongly correlated with the local backbone conformation (Fig. S4), such that the corresponding combinations are rare in the MD simulations. Thus the 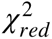 score can be effectively optimized by local optimization of the dipeptide unit, but the required conformations are difficult to recover through ensemble selection over the whole protein. These cases illustrate both the sensitivity of NOESY-based ensemble selection to local conformational detail and the limitations imposed by the sampled conformational space. We do not pursue them further here, since doing so would require deliberately increasing the population of rare conformational states rather than selecting from the equilibrium MD ensemble.

## DISCUSSION

In this work, we present the CoMAND2 platform for NMR structure determination of proteins via ensemble selection. Ensemble selection methods have been applied in NMR structure determination since at least the mid-1990s. At that time, their application to *de novo* structure determination was severely constrained by the need to explore conformational space. Since then, this conceptual framework has found a range of applications, from small molecules and peptides, where the search space is more tractable, to Bayesian reweighting schemes aimed at refining force-field parameters against experimental data. A notable example is provided by studies from the Al-Hashimi group, which used ensemble selection to characterize the internal dynamics of the HIV-1 transactivation response element (TAR) RNA ^34^. In these studies, large conformational pools were generated using RNA structure prediction tools and subsequently refined against extensive RDC data. These applications illustrate the power of ensemble selection in linking simulation and experiment. They also suggest that the primary limitations of the technique are not conceptual. Instead they reflect the dual challenges of adequately sampling around a suitable structural prior and obtaining quantitative data that can characterize deviations from that prior. In this context, the methods presented here greatly extend the applicability of ensemble selection. With the increasing availability of reliable structural models and efficient means of generating conformational diversity, sampling is no longer limiting. The ability to incorporate NOESY data as a quantitative observable adds a rich and widely accessible source of experimental information to drive the selection process. This provides access to the full range of folded protein targets studied by NMR.

As an example of the CoMAND2 method, we provide a new ensemble for human ubiquitin that faithfully recapitulates input data from a 3D CNH-NOESY experiment, together with complementary RDC and scalar coupling data. The accompanying residue-resolved χ² profile (Fig. 6A) provides a direct measure of how well the ensemble explains the experimental data and identifies regions where discrepancies remain. Thus, ensemble selection recasts structure determination as a statistical model-selection problem, placing quantitative validation at the centre of the structure determination process. In contrast, conventional NMR ensembles generated via restrained MD are only partially evaluated against the original observables. For NOESY data, validation typically relies on residual violations of the derived distance restraints. However, deriving restraints ultimately requires assigning NOESY cross-peaks to specific distances, a key interpretive step that separates the structural model from the underlying spectra. Under these conditions, residual violations primarily reflect the internal consistency of the restraint set. Yet this very consistency is the basic objective of NOESY assignment procedures, limiting the independence of restraint violations as a validation measure. As a result, ensemble selection approaches enable a more direct and quantitative form of validation against experimental data than has typically been available for conventional NMR ensembles.

Another difference between conventional structure determination and ensemble selection is the role of the forcefield. In conventional restrained MD, force constants on covalent geometry are usually kept very high to avoid distortion by the restraints. This can be seen as reducing the effective degrees of freedom of the structure determination process to the torsion angles around rotatable bonds; a relatively tractable problem, given the available data. In contrast, the choice of forcefield in ensemble selection is not directly constrained by the structure determination process. This independence is a key criterion for combining experiment and simulation. However, the forcefield does play a role, in that it acts as an implicit Bayesian prior; conformations with high energy will be rare in the conformer pool and can only be selected with concomitant support from the data. For NMR data this will mean that the capacity of the pool to explain the data will depend on accurately reproducing proton positions.Yet MD force fields are generally parameterized to reproduce molecular energetics and electrostatics rather than the effective hydrogen positions required to reproduce NMR observables. Such small deviations can nevertheless have a substantial effects, and improved RDC fits are obtainable through minor adjustments of proton positions ^35^. Considering the r^-6^ distance dependence of the NOE, NOESY data should be even more sensitive. A change in inter-proton distance of just 0.05 Å at a medium distance of ∼3 Å will result in an approximately 10% change in peak intensity. Accordingly, similar improvements in fit can be obtained for CNH-NOESY data, for example by adjustment of amide bond lengths via simple gradient minimization, and such methods are readily implemented in the CoMAND2 framework (Fig. S3). However, we do not currently have a sufficient basis for determining whether such refinements represent genuine conformational information rather than absorption of forcefield or forward-model error. Together with the above discussion of rare conformers in the hUb optimization, this highlights a growing interdependence between conformational sampling methods and the forcefields used to generate them. As ensemble-selection approaches increasingly probe low-population substates and long-range structural correlations, forcefields and selection methodologies may need to evolve in parallel to capture the full information content of NOESY data.

Figures 4 and 5 presents two examples of ensemble selection in bespoke state spaces, highlighting a major advantage of the current framework: the ability to focus conformational sampling on specific structural hypotheses. In this mode, the method conceptually aligns with Bayesian reweighting approaches, extending the application of NOESY data beyond conventional structure determination and toward targeted ensemble inference. The modular design of CoMAND2 should allow flexible integration of both NMR observables and complementary biophysical data in order to address specific mechanistic questions. Of particular interest will be the integration of structural and dynamics information, where conformational diversity can be correlated with motional timescales. This perspective aligns with recent ideas of 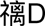 structure determination”, in which structural ensembles reflect not only conformational heterogeneity but also associated dynamic timescales. In this context, the CoMAND2 platform can form part of a toolkit of methods that shift the paradigm of NMR structure determination beyond restrained-MD, toward direct representations of protein conformational diversity.

## MATERIALS AND METHODS

### Forward Models

#### NOESY

In the original CoMAND implementation, we introduced SHINE (Simulating Hetero-Indirect NOESY Experiments) to back-calculate multidimensional NOESY spectra from molecular coordinates ^21^. SHINE was developed based on a previous literature implementation ^19^, with extensions enabling simulation of spectra with arbitrary combinations of ^1^H, ^13^C and ^15^N dimensions. To enable integration into the CoMAND2 API, we have refactored SHINE in Python, restricting its scope to the calculation of one-dimensional sub-spectra (“strips”) corresponding to contacts involving a single proton or proton group. These strips are directly comparable to vectors extracted from experimental spectra.

At the heart of NOESY back-calculation is the construction of the relaxation matrix **R** from internuclear distances. The off-diagonal elements of **R** are the cross-relaxation rates *σ_ij_* between spins *i* and *j*, which are computed from the internuclear distance *d_ij_* and spectral density functions derived from a motional model for the inter-proton vector. For the current application, we employ a reduced model based on a single global isotropic correlation time, supplemented by empirical corrections for fast internal motions such as methyl rotation. This provides a simplified motional description of NOE relaxation that is computationally efficient while retaining the dominant contributions relevant for back-calculation. The full relaxation matrix contains both homonuclear proton–proton terms and heteronuclear contributions involving isotope-labelled heavy atoms. Because heteronuclear relaxation is dominated by local covalent geometry, in particular the directly bonded heavy atom, these contributions can be approximated using atom-level class averages. This reduces **R** to its proton–proton block, significantly reducing computational cost with minimal impact on the back-calculated observables.

For a single, static structure, the matrix of expected NOESY intensities **I**, can be calculated from **R** as:

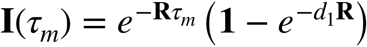

where *τ_m_* is the mixing time of the experiment and *d*_1_ the relaxation delay. The most expensive element of this calculation is the exponentiation of **R**, which can be performed with an efficient algorithm based on spectral decomposition, valid for any function *f* defined on the eigenvalues of **R**:

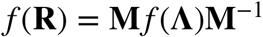

where **M** is a matrix of the eigenvectors of **R** and **Λ** is the diagonal matrix of eigenvalues. As **R** is a real, symmetric matrix, **M** can be obtained with a symmetric eigensolver, and **M**^−1^ = **M***^T^*. The exponential is then:

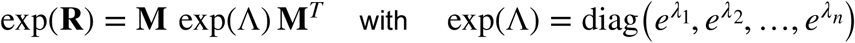

As this algorithm scales as 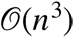 with matrix size (i.e. the number of protons) we apply a fragmentation approach, constructing **R** from atoms within a shell around an atom of interest. This shell contains all residues contacting the central residue based on pre-calculated residue radii, plus an adjustable buffer. Typically, back-calculated intensities converge as this buffer approaches 2 Å, and fragments contain 100-150 atoms.

To build the back-calculated strips, the intensity matrix is first filtered to extract contacts relevant to the atom of interest and spectrum type. Each intensity is represented as a Gaussian multiplet in the back-calculated vector, defined by user-specified chemical shifts and line widths and accounting for folding of peaks outside the spectral window. The primary purpose of these strips is quantitative comparison with experimental data, and mismatches in chemical shifts or line widths directly impact the minimum achievable residual. These parameters are therefore calibrated against individual experimental lines prior to any optimization, lowering this residual floor without introducing additional degrees of freedom into the global fit. We have also implemented a correction for the ^13^C excitation profile, as the relatively wide spectral width in the ^13^C dimension can lead to systematic attenuation of signals far from the spectral centre, in particular aromatic resonances. Remaining intensity-modulating effects primarily affect the relative scaling between strips and are incorporated into a global scaling factor in the fitting procedure.

#### Couplings

Back-calculation of RDCs from molecular coordinates is straightforward using the Saupe matrix formulation of the alignment tensor describing the orientational distribution of the medium:

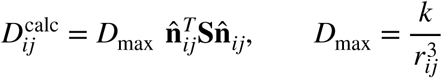

where *D*_max_ is the maximum coupling for a pair of spins *i* and *j*, **^n̂^***_ij_* is the unit vector along the internuclear axis, *r_ij_* the internuclear distance and *k* collects the relevant physical constants. The Saupe matrix, **S,** is a symmetric, traceless, 3×3 tensor, where the traceless condition (Tr(**S**) = 0), reflects the zero isotropic average of dipolar couplings and implies that **S** has five independent parameters. For computational convenience, the five independent components of the Saupe tensor are rearranged into the vector:

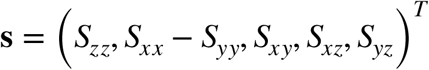

such that:

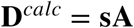

where **A** is the design matrix containing geometric coefficients derived from the internuclear vectors. In this form, the alignment tensor can be obtained directly from experimental RDCs and reference coordinates using linear least squares.

The current implementation provides two methods for RDC back-calculation: a fitted-tensor approach, in which the alignment tensor is determined independently for each structure, and a fixed-tensor approach, in which a predefined alignment tensor is used. In the latter case, RDC back-calculation is performed by rotating the molecular coordinates into the principal frame of **S** and evaluating the corresponding internuclear vectors. Both approaches employ the same tensor formalism. The fitted-tensor method treats the five independent tensor parameters as global variables, whereas the fixed-tensor method constrains the tensor magnitude and rhombicity, fitting only the three Euler angles describing the orientation between the tensor and molecular frames. The corresponding RDC fits therefore consume five and three degrees of freedom, respectively. Internuclear distances may be obtained directly from atomic coordinates or assigned fixed values. Fixed values based on the parameterization of Ottiger and Bax ^36^ are also provided. This ensures that *D*_max_ can be defined consistently for tensors determined by external programs.

For back-calculation of scalar couplings, we employ parameterized relationships between coupling constants and local geometry, as these are computationally tractable for larger pools of structures. Vicinal couplings are described using Karplus-type relationships between coupling constants and torsion angles. The current implementation allows arbitrary couplings to be defined through user-supplied Karplus parameters. Default parameter sets are also provided for commonly observed couplings using parameterizations derived from quantum-chemical calculations ^37^. Hydrogen-bond mediated couplings are treated analogously following the formulation of Barfield ^33^. User-defined parameters may be supplied for a two-parameter model relating the coupling to hydrogen-bond geometry:

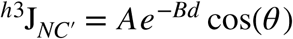

where *d* is the acceptor-donor distance and *θ* the acceptor antecedent angle. For backbone hydrogen bonds, a default parameterization is also provided, including empirical corrections for secondary structure.

### Scoring

#### The χ² scoring function

To provide a scoring function capable of handling multiple data types, we employ a χ²-based metric in which deviations between back-calculated and experimental values for measurement *i* are normalized by the corresponding estimate of the experimental uncertainty, 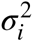:

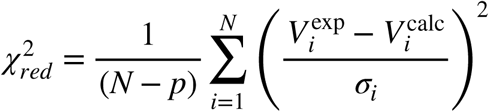

The reduced χ²accounts for the effective number of degrees of freedom, with *N* the the number of observations and *p* the number of fitted parameters. The definition of both the error estimates and the number of fitted parameters depends on the data type and fitting mode.

For NOESY data, a global scaling factor *s* is introduced to account for arbitrary intensity normalization between strips. The optimal value of *s* is obtained analytically by minimization of the residual ^21^ and applied prior to evaluation of the scoring function, *V* ^calc^ → *sV* ^calc^. This introduces one fitted parameter to the effective degrees of freedom. For overlapped strips, deconvolution is performed via weighting factors, as described previously ^21^ introducing w咵additional fitted parameters, where w is the number of strips contributing to the overlap. Estimation of the uncertainty for NOESY-derived observables requires a model of the intensity-dependent noise across experimental strips. We therefore employ a heteroscedastic noise model consisting of a baseline component and a signal-dependent component. To capture the approximately linear scaling of noise amplitude with signal intensity, the noise variance is modeled as:

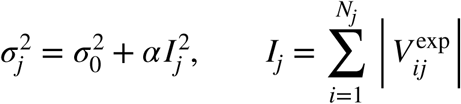

where *I_j_* is the sum of the absolute intensity for strip *j*.

The baseline noise variance, 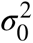 is estimated empirically by sampling strips from regions of the multidimensional spectrum that are well separated from known signal positions. To reduce contamination from residual signals and artifacts, candidate noise strips are filtered by rejecting those containing intensities exceeding a fixed threshold (typically 10σrelative to the initial noise estimate). The selected strips are used to parameterize a multivariate Gaussian model describing the baseline noise distribution. To account for the finite bandwidth of indirect dimensions, the model is constrained using a second-order Butterworth low-pass filter with a cutoff corresponding to half the spectral sweep width (Hz), ensuring consistency with the effective frequency content of the experiment. Synthetic noise vectors drawn from this distribution are then used to estimate the variance. The signal-dependent component of the model accounts for error sources not captured by sampling of empty regions, with the magnitude of this contribution controlled by a single global parameter, α. In the limit of high strip intensity, the signal-dependent term dominates the uncertainty estimate and αtherefore represents the asymptotic fractional uncertainty. The value of αis estimated from the dependence of the overall variance of residuals on the square of strip intensity and based on a selection of high-quality fits covering the intensity range of the experiment (Fig. S1).

For scalar couplings, the Karplus parameters are fixed *a priori* and no parameters are fitted to the experimental data, such that the degrees of freedom for the 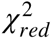 calculation are equal to the number of observed couplings. For RDC data, the number of fitted parameters depends on the tensor fitting mode, with the fitted- and fixed-tensor approaches consuming five and three degrees of freedom, respectively. For both data types, error estimation is performed by iterative global fitting, initialized using the reported experimental uncertainty, followed by linear regression of experimental against back-calculated values. The standard deviation is then updated from the regression and the procedure repeated until convergence. The resulting effective standard deviations are used in the calculation of 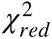. For RDCs, global fitting is performed over all couplings for each alignment tensor, while standard deviations are estimated separately for each coupling type.

#### Regularization

The χ²-based scoring function described above provides a measure of agreement with experimental data across multiple data types. To prevent overfitting during optimization, it is augmented with a regularization term based on the effective number of states, N_eff_, allowing control over ensemble complexity.

For an ensemble containing N_ens_ conformers and a state space comprising M states, the effective number of states is calculated as:

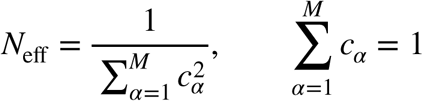

where c_α_ denotes the normalized occupancy of state α, defined as the fraction of ensemble members assigned to that state:

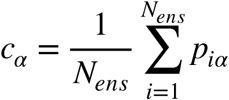

where *p_iα_* is the denotes the membership weight of conformer *i* in state α. To provide a flexible regularization framework, the current implementation allows memberships to be defined in arbitrary state spaces assigned to individual residues. For hard membership, the state space is partitioned into discrete bins and occupancies are calculated from the resulting normalized bin counts. For soft membership, Gaussian mixture models (GMMs) are trained on data that sample the state space, for example structural databases or molecular dynamics trajectories. To define default soft regularization spaces, this procedure was applied to data from the MolProbity database ^38^. For each residue type, a dihedral space was defined comprising ψ_r-1_, ω_r-1_, φ_r_ and up to two side-chain dihedrals (χ_1_, χ_2_).The optimum number of clusters was first determined using k-means clustering, with the results evaluated using the Akaike Information Criterion. GMMs were then trained for each residue type using the resulting cluster centers as starting points. The resulting GMM parameters are provided in portable JSON format with the CoMAND2 source code.

To combine the N_eff_-based regularization term with the χ^2^ scoring function we apply a linear combination:

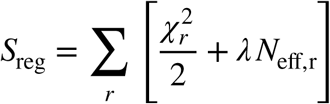

where λis a scaling factor controlling the trade-off between fit quality and ensemble complexity. Both 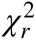 and *N*_e,r_ are evaluated independently within each regularization space, *r* by defining 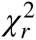 over the subset of observables that report on that space. This ensures consistency with the construction of the membership spaces. The resulting function is used as an optimization objective and is not intended to represent a probabilistic or generative model.

#### Ensemble Selection

In the ensemble selection procedure, forward models are used to compute predicted experimental observables for each conformer in the conformer pool. These predictions are assembled into structured data matrices, providing a mapping between individual conformers and the corresponding experimental observables. In a corresponding manner, features defining the regularization space are mapped onto conformers to construct membership matrices. The scoring function is then evaluated as a weighted combination of these matrices under a set of ensemble weights. Ensemble selection is performed by setting the ensemble weights - and thus the ensemble composition - to minimize the objective function.

The current implementation provides a Monte-Carlo simulated annealing scheme (MCSA) for ensemble selection that is robust to potentially rugged scoring landscapes. The heart of this algorithm is sampling of ensemble compositions via random changes to a starting ensemble. At each iteration, a conformer is exchanged between the ensemble and conformer pool to generate a trial ensemble. Trial ensembles that improve the score are accepted unconditionally, while those that worsen the score are accepted with probability:

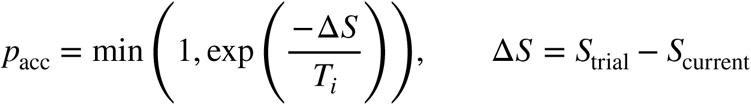

where *T_i_* is the pseudo-temperature at iteration *i*. The characteristics of the algorithm are governed largely by the temperature profile, with current implementation supporting two different modes. In the linear ramp mode, the temperature is reduced at each iteration at a constant rate. In adaptive mode the temperature is only reduced after accepted moves, allowing the algorithm to dedicate more time to exploring flat regions of the scoring landscape. In the default protocol we combine these modes in a two-stage strategy with a fixed number of iterations at high temperature, followed by adaptive cooling until convergence. Typically, the regularization term is disabled during the first stage (λ_Neff_ = 0), allowing a broader exploration of solution space before regularization is introduced during cooling. Convergence can be controlled using both the total number of iterations and the number of iterations performed at the current temperature. In practice, repeated runs from different random starting ensembles yield highly reproducible final scores for conformer pools containing up to 40,000 structures.

The regularization parameter, λ_Neff_ is estimated empirically from the typical improvement in fit quality associated with an increase in ensemble complexity. Starting from a small ensemble, random perturbations are generated by adding a single conformer. Perturbations that both improve the fit and increase N_eff_ are retained. λ_Neff_ is then calculated as:

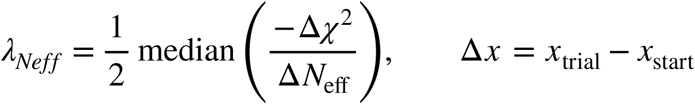

This captures the characteristic trade-off between improved agreement with experimental data and increased ensemble complexity.

#### Regularization of (P3-7)_3_

To regularize the single α-helical peptide (P3-7)3 ^30^ we employed the same pool of 26,000 MD frames used in the original study ^31^, back-calculating CNH-NOESY data for each frame using the current version of the program. These were compared with the experimental CNH-NOESY data from the original study. For each glutamine residue, conformational membership was defined using rotamer bins centered at -60°, 180° and 60° for both the χ_1_ and χ_2_ dihedral angles, with a bin width of 30°. Dihedrals falling outside these three bins were assigned to a fourth, non-rotameric category, giving 16 possible χ_1_/χ_2_ combinations for each residue. Hard membership was then represented by a binary vector for each member of the conformer pool, with 1 indicating membership and 0 otherwise. Compiling these vectors into a matrix allows the overall membership of an ensemble to be calculated conveniently as the matrix–weight product using the ensemble weights obtained during MCSA optimization.

To examine the response of the system to increasing regularization pressure, we performed MCSA optimization at five values of λ_Neff_ ranging from 0 to 1, while simultaneously optimizing agreement with CNH-NOESY data for the backbone amide and side-chain HE21 protons of each glutamine residue, together with the amide proton of the following residue. These provide sensitive reporters of side-chain conformation. At each value of λ_Neff_, 50 ensembles of 12 structures were generated from random starting ensembles. Data for Figures 5A and 5B were taken from the best-scoring ensemble at each λ_Neff_ value, whereas the histogram in Figure 5C was compiled from all structures occurring in the 50 ensembles at each value, weighted by their frequency of use.

#### Structure Determination of human Ubiquitin

To construct the updated ensemble for hUb, we used the same pool of 40,000 frames generated by equilibrium MD in explicit solvent as employed in the original study ^21^. CNH-NOESY data for each frame were back-calculated using the current version of the program. The experimental CNH-NOESY data from the original study were used, complemented by the four sets of RDC data,^3^J_HNHα_ and ^h3^J_NC’_ data used by Maltsev et al. (2MJB) ^32^. H_α_–C_α_ and side-chain C–H RDCs were omitted because these measurements are typically obtained from protonated samples, whereas the other RDCs may be obtained from deuterated samples and would therefore require separate treatment of the alignment tensor. Regularization was performed using the default soft assignment spaces, with membership vectors constructed for each residue based on residue type for each structure in the pool.

To select the initial ensemble, MCSA selection was performed by optimizing over all CNH-NOESY, RDC and J-coupling data. To establish an appropriate value of λ_Neff_, an initial unregularized optimization was performed, and the slope of the regularization curve near this solution was probed by random perturbations. A value of λ_Neff_ = 0.2 was selected. Twenty independent ensembles of 12 members were then selected from random starting ensembles, and the ensemble with the best S_reg_ score chosen for further analysis.

In addition, individual optimizations were performed for dipeptide fragments, co-optimizing the contributions from residues i and i+1. These provided a baseline S_reg_ score for each residue. Residues in the selected ensemble with scores substantially above this baseline were considered for manual rebuilding to alter side-chain rotamer distributions using ISOLDE ^39^.The effect of these changes was monitored using the local 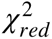 score. Each member of the modified ensemble was then used to seed an independent MD simulation using the same protocol used to generate the original conformer pool ^21^, generating approximately 500 frames per structure (4,960 frames in total). A final round of regularized selection was performed on this smaller pool using all experimental data, yielding the final ensemble.

## References

(1) Bowman, G. R. AlphaFold and Protein Folding: Not Dead Yet! The Frontier Is Conformational Ensembles. Annual Review of Biomedical Data Science 2024, 7 (1), 51–57. 10.1146/annurev-biodatasci-102423-011435.

(2) Laurents, D. V. AlphaFold 2 and NMR Spectroscopy: Partners to Understand Protein Structure, Dynamics and Function. Front. Mol. Biosci. 2022, 9, 906437. 10.3389/fmolb.2022.906437.

(3) Rennie, M. L.; Oliver, M. R. Emerging Frontiers in Protein Structure Prediction Following the AlphaFold Revolution. J. R. Soc. Interface. 2025, 22 (225), 20240886. 10.1098/rsif.2024.0886.

(4) Mittermaier, A.; Kay, L. E. New Tools Provide New Insights in NMR Studies of Protein Dynamics. Science 2006, 312 (5771), 224–228. 10.1126/science.1124964.

(5) Henzler-Wildman, K.; Kern, D. Dynamic Personalities of Proteins. Nature 2007, 450 (7172), 964–972. 10.1038/nature06522.

(6) Palmer, A. G. Enzyme Dynamics from NMR Spectroscopy. Acc Chem Res 2015, 48 (2), 457–465. 10.1021/ar500340a.

(7) Lindorff-Larsen, K.; Best, R. B.; DePristo, M. A.; Dobson, C. M.; Vendruscolo, M. Simultaneous Determination of Protein Structure and Dynamics. Nature 2005, 433 (7022), 128– 132. 10.1038/nature03199.

(8) Bonomi, M.; Heller, G. T.; Camilloni, C.; Vendruscolo, M. Principles of Protein Structural Ensemble Determination. Current Opinion in Structural Biology 2017, 42, 106–116. 10.1016/j.sbi.2016.12.004.

(9) Rieping, W.; Habeck, M.; Bardiaux, B.; Bernard, A.; Malliavin, T. E.; Nilges, M. ARIA2: Automated NOE Assignment and Data Integration in NMR Structure Calculation. Bioinformatics 2007, 23 (3), 381–382. 10.1093/bioinformatics/btl589.

(10) Würz, J. M.; Kazemi, S.; Schmidt, E.; Bagaria, A.; Güntert, P. NMR-Based Automated Protein Structure Determination. Archives of Biochemistry and Biophysics 2017, 628, 24–32. 10.1016/j.abb.2017.02.011.

(11) Jumper, J.; Evans, R.; Pritzel, A.; Green, T.; Figurnov, M.; Ronneberger, O.; Tunyasuvunakool, K.; Bates, R.; Žídek, A.; Potapenko, A.; Bridgland, A.; Meyer, C.; Kohl, S. A. A.; Ballard, A. J.; Cowie, A.; Romera-Paredes, B.; Nikolov, S.; Jain, R.; Adler, J.; Back, T.; Petersen, S.; Reiman, D.; Clancy, E.; Zielinski, M.; Steinegger, M.; Pacholska, M.; Berghammer, T.; Bodenstein, S.; Silver, D.; Vinyals, O.; Senior, A. W.; Kavukcuoglu, K.; Kohli, P.; Hassabis, D. Highly Accurate Protein Structure Prediction with AlphaFold. Nature 2021, 596 (7873), 583–589. 10.1038/s41586-021-03819-2.

(12) Jänes, J.; Beltrao, P. Deep Learning for Protein Structure Prediction and Design—Progress and Applications. Mol Syst Biol 2024, 20 (3), 162–169. 10.1038/s44320-024-00016-x.

(13) Rieping, W.; Habeck, M.; Nilges, M. Inferential Structure Determination. Science 2005, 309 (5732), 303–306. 10.1126/science.1110428.

(14) Bothe, J. R.; Nikolova, E. N.; Eichhorn, C. D.; Chugh, J.; Hansen, A. L.; Al-Hashimi, H. M. Characterizing RNA Dynamics at Atomic Resolution Using Solution-State NMR Spectroscopy. Nat Methods 2011, 8 (11), 919–931. 10.1038/nmeth.1735.

(15) Salmon, L.; Bascom, G.; Andricioaei, I.; Al-Hashimi, H. M. A General Method for Constructing Atomic-Resolution RNA Ensembles Using NMR Residual Dipolar Couplings: The Basis for Interhelical Motions Revealed. J. Am. Chem. Soc. 2013, 135 (14), 5457–5466. 10.1021/ja400920w.

(16) Hummer, G.; Köfinger, J. Bayesian Ensemble Refinement by Replica Simulations and Reweighting. J Chem Phys 2015, 143 (24), 243150. 10.1063/1.4937786.

(17) Chen, Y.; Campbell, S. L.; Dokholyan, N. V. Deciphering Protein Dynamics from NMR Data Using Explicit Structure Sampling and Selection. Biophysical Journal 2007, 93 (7), 2300–2306. 10.1529/biophysj.107.104174.

(18) Gippert, G. P.; Yip, P. F.; Wright, P. E.; Case, D. A. Computational Methods for Determining Protein Structures from NMR Data. Biochemical Pharmacology 1990, 40 (1), 15–22. 10.1016/0006-2952(90)90172-H.

(19) Zhu, L.; Dyson, H. J.; Wright, P. E. A NOESY-HSQC Simulation Program, SPIRIT. J Biomol NMR 1998, 11 (1), 17–29. 10.1023/a:1008252526537.

(20) Diercks, T.; Coles, M.; Kessler, H. An Efficient Strategy for Assignment of Cross-Peaks in 3D Heteronuclear NOESY Experiments. J Biomol NMR 1999, 15 (2), 177–180. 10.1023/a:1008367912535.

(21) ElGamacy, M.; Riss, M.; Zhu, H.; Truffault, V.; Coles, M. Mapping Local Conformational Landscapes of Proteins in Solution. Structure 2019, 27 (5), 853–865.e5. 10.1016/j.str.2019.03.005.

(22) Tolman, J. R.; Flanagan, J. M.; Kennedy, M. A.; Prestegard, J. H. Nuclear Magnetic Dipole Interactions in Field-Oriented Proteins: Information for Structure Determination in Solution. Proc Natl Acad Sci U S A 1995, 92 (20), 9279–9283. 10.1073/pnas.92.20.9279.

(23) Tjandra, N.; Bax, A. Direct Measurement of Distances and Angles in Biomolecules by NMR in a Dilute Liquid Crystalline Medium. Science 1997, 278 (5340), 1111–1114. 10.1126/science.278.5340.1111.

(24) Eastman, P.; Galvelis, R.; Peláez, R. P.; Abreu, C. R. A.; Farr, S. E.; Gallicchio, E.; Gorenko, A.; Henry, M. M.; Hu, F.; Huang, J.; Krämer, A.; Michel, J.; Mitchell, J. A.; Pande, V. S.; Rodrigues, J. P.; Rodriguez-Guerra, J.; Simmonett, A. C.; Singh, S.; Swails, J.; Turner, P.; Wang, Y.; Zhang, I.; Chodera, J. D.; De Fabritiis, G.; Markland, T. E. OpenMM 8: Molecular Dynamics Simulation with Machine Learning Potentials. J Phys Chem B 2024, 128 (1), 109–116. 10.1021/acs.jpcb.3c06662.

(25) Cornilescu, G.; Bax, A. Measurement of Proton, Nitrogen, and Carbonyl Chemical Shielding Anisotropies in a Protein Dissolved in a Dilute Liquid Crystalline Phase. J. Am. Chem. Soc. 2000, 122 (41), 10143–10154. 10.1021/ja0016194.

(26) Hill, M. O. Diversity and Evenness: A Unifying Notation and Its Consequences. Ecology 1973, 54 (2), 427–432. 10.2307/1934352.

(27) Jost, L. Entropy and Diversity. Oikos 2006, 113 (2), 363–375. 10.1111/j.2006.0030-1299.14714.x.

(28) Shi, H.; Rangadurai, A.; Abou Assi, H.; Roy, R.; Case, D. A.; Herschlag, D.; Yesselman, J. D.; Al-Hashimi, H. M. Rapid and Accurate Determination of Atomistic RNA Dynamic Ensemble Models Using NMR and Structure Prediction. Nat Commun 2020, 11 (1), 5531. 10.1038/s41467-020-19371-y.

(29) Korycinski, M.; Albrecht, R.; Ursinus, A.; Hartmann, M. D.; Coles, M.; Martin, J.; Dunin-Horkawicz, S.; Lupas, A. N. STAC--A New Domain Associated with Transmembrane Solute Transport and Two-Component Signal Transduction Systems. J Mol Biol 2015, 427 (20), 3327–3339. 10.1016/j.jmb.2015.08.017.

(30) Escobedo, A.; Topal, B.; Kunze, M. B. A.; Aranda, J.; Chiesa, G.; Mungianu, D.; Bernardo-Seisdedos, G.; Eftekharzadeh, B.; Gairí, M.; Pierattelli, R.; Felli, I. C.; Diercks, T.; Millet, O.; García, J.; Orozco, M.; Crehuet, R.; Lindorff-Larsen, K.; Salvatella, X. Side Chain to Main Chain Hydrogen Bonds Stabilize a Polyglutamine Helix in a Transcription Factor. Nat Commun 2019, 10 (1), 2034. 10.1038/s41467-019-09923-2.

(31) Escobedo, A.; Piccirillo, J.; Aranda, J.; Diercks, T.; Mateos, B.; Garcia-Cabau, C.; Sánchez-Navarro, M.; Topal, B.; Biesaga, M.; Staby, L.; Kragelund, B. B.; García, J.; Millet, O.; Orozco, M.; Coles, M.; Crehuet, R.; Salvatella, X. A Glutamine-Based Single α-Helix Scaffold to Target Globular Proteins. Nat Commun 2022, 13 (1), 7073. 10.1038/s41467-022-34793-6.

(32) Maltsev, A. S.; Grishaev, A.; Roche, J.; Zasloff, M.; Bax, A. Improved Cross Validation of a Static Ubiquitin Structure Derived from High Precision Residual Dipolar Couplings Measured in a Drug-Based Liquid Crystalline Phase. J. Am. Chem. Soc. 2014, 136 (10), 3752–3755. 10.1021/ja4132642.

(33) Barfield, M. Structural Dependencies of Interresidue Scalar Coupling^h3^ JNC‘ and Donor1 H Chemical Shifts in the Hydrogen Bonding Regions of Proteins. J. Am. Chem. Soc. 2002, 124 (15), 4158–4168. 10.1021/ja012674v.

(34) Liu, B.; Shi, H.; Al-Hashimi, H. M. Developments in Solution-State NMR Yield Broader and Deeper Views of the Dynamic Ensembles of Nucleic Acids. Current Opinion in Structural Biology 2021, 70, 16–25. 10.1016/j.sbi.2021.02.007.

(35) Grishaev, A.; Ying, J.; Bax, A. Imino Hydrogen Positions in Nucleic Acids from Density Functional Theory Validated by NMR Residual Dipolar Couplings. J. Am. Chem. Soc. 2012, 134 (16), 6956–6959. 10.1021/ja301775j.

(36) Ottiger, M.; Bax, A. Determination of Relative N−H^N^, N−C‘, Cα −C‘, and Cα −Hα Effective Bond Lengths in a Protein by NMR in a Dilute Liquid Crystalline Phase. J. Am. Chem. Soc. 1998, 120 (47), 12334–12341. 10.1021/ja9826791.

(37) Vögeli, B.; Ying, J.; Grishaev, A.; Bax, A. Limits on Variations in Protein Backbone Dynamics from Precise Measurements of Scalar Couplings. J. Am. Chem. Soc. 2007, 129 (30), 9377–9385. 10.1021/ja070324o.

(38) Williams, C. J.; Headd, J. J.; Moriarty, N. W.; Prisant, M. G.; Videau, L. L.; Deis, L. N.; Verma, V.; Keedy, D. A.; Hintze, B. J.; Chen, V. B.; Jain, S.; Lewis, S. M.; Arendall, W. B.; Snoeyink, J.; Adams, P. D.; Lovell, S. C.; Richardson, J. S.; Richardson, D. C. MolProbity: More and Better Reference Data for Improved All-atom Structure Validation. Protein Science 2018, 27 (1), 293–315. 10.1002/pro.3330.

(39) Croll, T. I. ISOLDE : A Physically Realistic Environment for Model Building into Low-Resolution Electron-Density Maps. Acta Crystallogr D Struct Biol 2018, 74 (6), 519–530. 10.1107/S2059798318002425.

(40) Müller, T. A.; Stahlecker, F.; Roganowicz, K.; Samir, S.; Weiss, G. L.; Coles, M.; Selim, K. A. The Calcium-Binding Protein CSE Links Ca2+ Signaling with Cell-Cell Communication in Cyanobacteria. EMBO J 2026. 10.1038/s44318-026-00893-y.

