## Supplemental Material for "Inferring protein ensembles directly from NOESY spectra"

### SUPPLEMENTARY MATERIAL

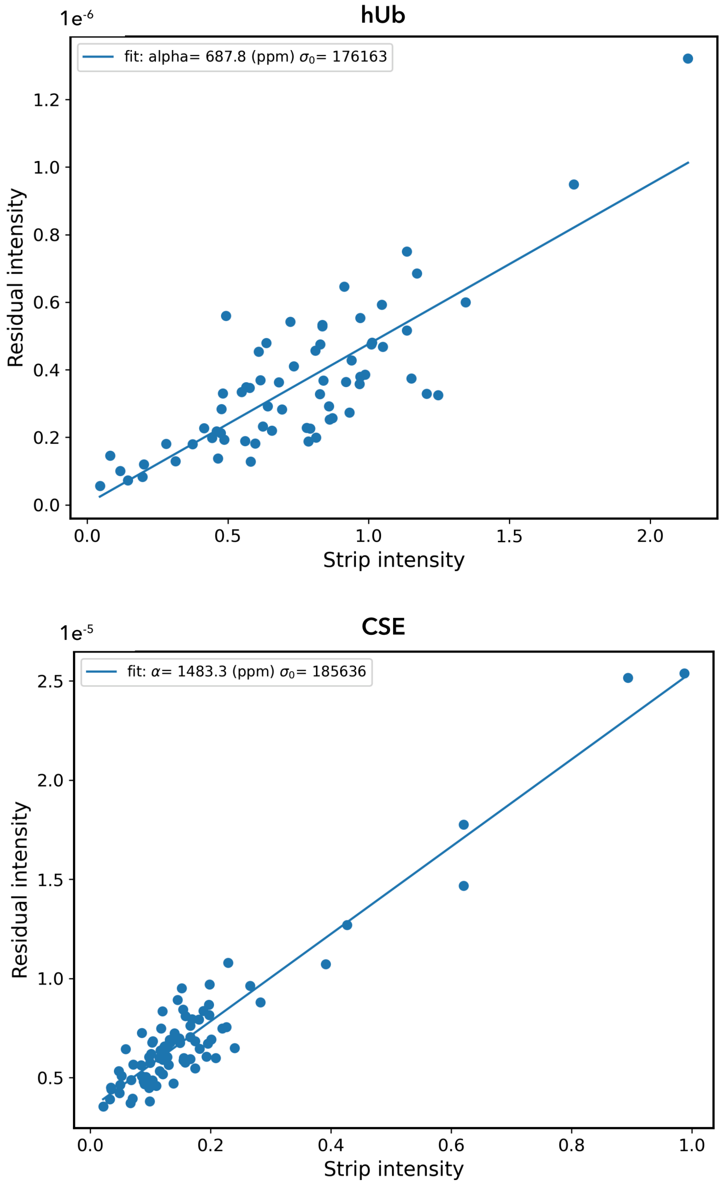


**Figure S1. The heteroscedastic noise model for NOESY data**

The plot shows the relationship between total strip intensity (arbitrary units) and the residual for a series of ensemble selection fits for human Ubiquitin (top) and CSE {Müller, 2026} (bottom). The fits are over a dipeptide (residue i and residue i+1) for all residues in the proteins. These fits typically produce low residuals (R-factor = 0.08 ± 0.02 and 0.28 ± 0.1, respectively), such that this residual is dominated by noise. Note that the residual axes are scaled by a factor of 1e$^{-6}$ relative to the total intensity. The slope of the fitted line provides the square of the parameter α in the heteroscedastic noise model:

$$\sigma_j^2 = \sigma_0^2 + \alpha I_j^2$$

where $\sigma_0^2$ is the baseline (thermal) noise and $\alpha I_j^2$ accounts for sources of error proportional to the signal intensity, such as $t_1$ noise, plus forward modeling error.

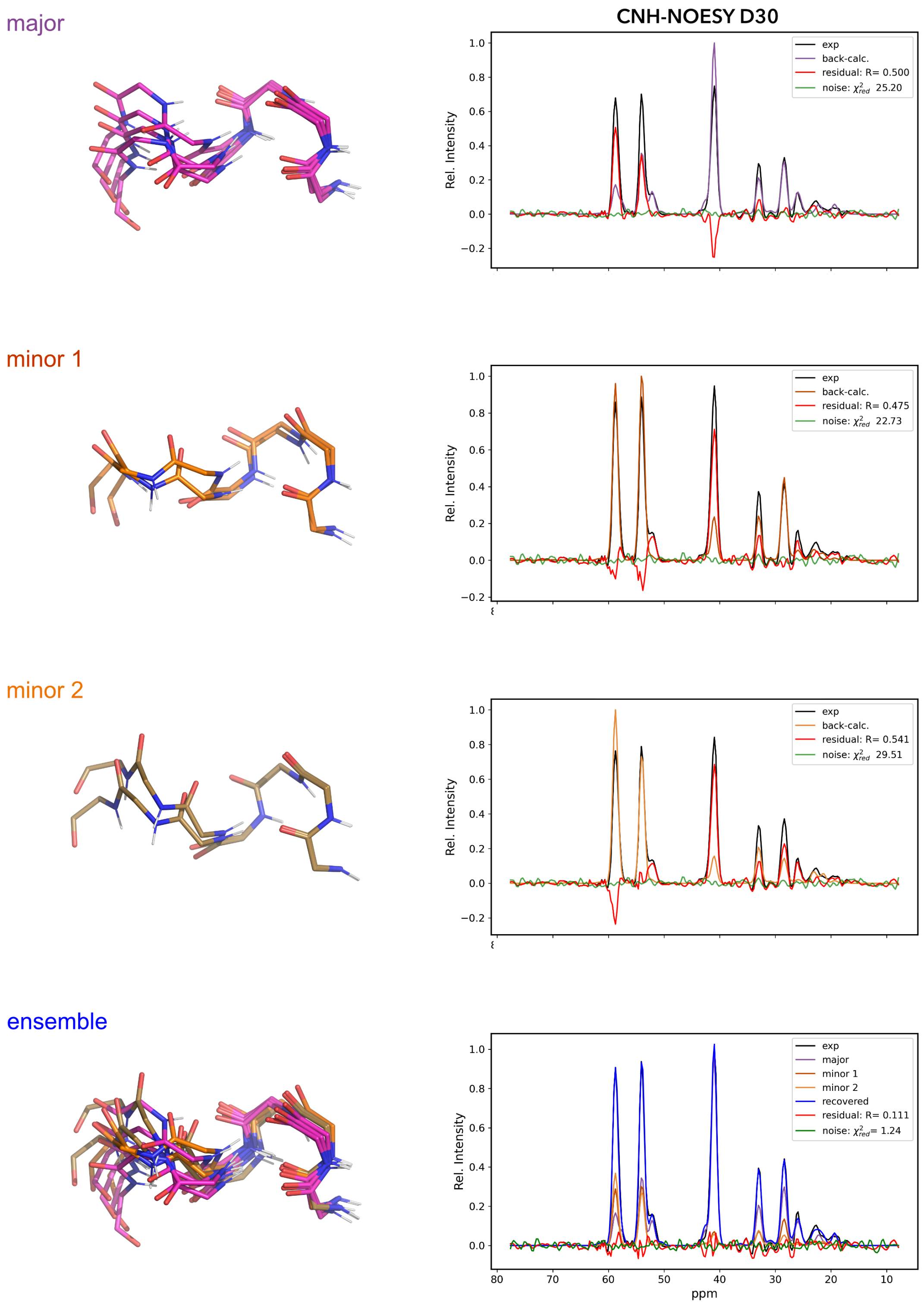


**Figure S2. A backbone polymorphism in Af1503 STAC**

Representative structures are shown for the backbone conformational states selected for D30 in Af1503 STAC {Korycinski, 2015}. The major conformation (62.5%) is shown in purple, while two minor conformations (~19% each) are in dark and light orange. For each conformer, a comparison between the back-calculated and experimental CNH-NOESY data is shown, with the back-calculated strip colored according to the conformer, the experimental data in black and the residual in red. For each comparison, the $\chi^2_{red}$ score is shown on the plot, based on the heteroscedastic noise model (baseline noise; green). No single conformer explains the experimental data well, whereas a linear combination of the three explains the data at close to the noise level.

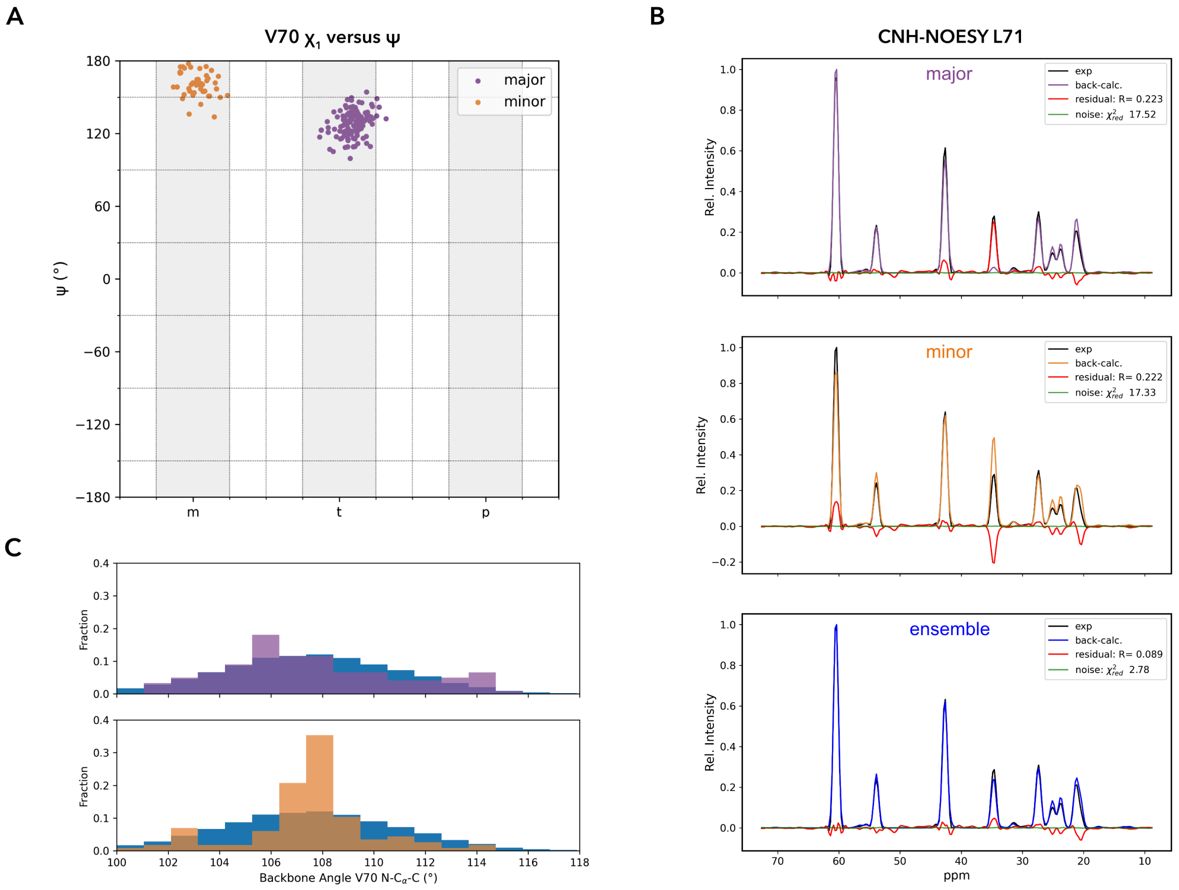


**Figure S4. Backbone/side chain correlation in human Ubiquitin**

The CNH-NOESY strip of L71 is sensitive to the conformation of V70, which shows a strong correlation between is backbone ψ and side chain $\chi_1$ angles. Data is show for 20 MCSA runs with random starting ensembles, co-optimizing over all data for V70 and L71. (A) Plot of ψ versus $\chi_1$ for V70, showing that the major and minor $\chi_1$ conformers segregate into distinct ψ clusters. (B) Both the major and minor conformers identified in panel A are required to explain the CNH-NOESY data. CNH-NOESY comparisons are shown, colored as in Fig. S2. (C) Histograms of the backbone N-C$_\alpha$-C' angle for V70 shown separately for the major and minor conformers against that for the complete conformer pool (blue). The minor conformer is restricted to a much narrower region of the background distribution, indicating that its formation depends on a more specific local backbone geometry.
